# Small molecule activation of m6A mRNA methylation as a novel approach for neuroprotection

**DOI:** 10.1101/2023.07.05.547860

**Authors:** Li-Ying Yu, Simona Selberg, Indrek Teino, Jinhan Nam, Larisa Ivanova, Brunaldo Renzi, Neinar Seli, Esko Kankuri, Merja H. Voutilainen, Mati Karelson, Mart Saarma

**Affiliations:** Institute of Biotechnology, HiLIFE, Viikinkaari 5D, Helsinki FIN-00014, University of Helsinki, Helsinki, Finland; Institute of Chemistry, University of Tartu, Ravila 14a, Tartu 50411, Estonia; Institute of Molecular and Cell Biology, University of Tartu, Riia 23, 50107, Estonia; Division of Pharmacology and Drug Therapy, Faculty of Pharmacy, University of Helsinki, Helsinki, Viikinkaari 5, FIN-00014, Finland; Chemestmed, Ltd., Riia 130b/2, Tartu 50411, Estonia; Faculty of Medicine, Department of Pharmacology, Helsinki FIN-00014, University of Helsinki, Finland

## Abstract

N6-Methyladenosine (m6A) is the most common mRNA base modification in eukaryotes. Methylation of adenosine residues to m6A contributes to the regulation of splicing, transport, stability, and translation of mRNA and two main classes of enzymes regulate it. The formation of m6A is catalysed by a methyltransferase complex containing methyltransferase-like 3 (METTL3), METTL14, and Wilms’ tumour 1-associated protein (WTAP) as well as monomeric METTL16. Demethylation of m6A is catalysed by the fat mass and obesity-associated protein FTO and the RNA demethylase AlkB homolog 5 (ALKBH5). The m6A mRNA methylation dysregulation occurs in the nervous system and in Parkinson’s disease (PD), but it remains poorly studied. Moreover, the role of m6A mRNA methylation in neuronal survival, neuroprotection, and neuroregeneration is unclear. We have earlier used high-throughput virtual screening of large compound libraries and identified four unique small-molecule ligands that activate m6A mRNA methylation by binding to the METTL3/14/WTAP complex and enhancing the binding of the methylation substrate SAM to nanomolar concentrations. Following this, we now discovered that two methyltransferase activators at 10 nM concentrations supported the survival and protected dopamine (DA) neurons in culture in growth factor deprivation and 6-hydroxydopamine (6-OHDA) neurotoxin models. In contrast, METTL3/14 inhibitor STM2457 triggered death of DA neurons. For clinical translation we also tested the most efficient compound C4 on induced pluripotent stem cell-derived human DA neurons and in animal model of Parkinson’s disease (PD). C4 compound protected human DA neurons from 6-OHDA-induced cell death and increased neurite outgrowth and the number of processes demonstrating that it has both neuroprotective and neurorestorative properties. METTL3/14 activator C4 improved motor behaviour and protected DA neurons and their fibres faster and much more efficiently than GDNF in the rat 6-OHDA model of PD. These are the first specific activators of METTL3/14/WTAP and first demonstration that m6A regulators can protect and regenerate neurons. These data demonstrate that m6A mRNA methylation is a novel pathway regulating neuronal survival and regeneration.

## Introduction

Chemical modifications of RNA bases have been shown to have critical impact on many cellular functions, such as proliferation, survival, and differentiation, mostly through regulation of RNA stability and transport (Anreiter et al., 2021; Tzelepis et al., 2019). The most abundant modified base in eukaryotic messenger RNA is N6-methyladenosine (m6A) that affects mRNA splicing, ribosomal translation, intracellular transport, and stability, thus playing thus a crucial role in regulating cell differentiation, cellular signalling, carcinogenesis, and immune tolerance (Maity & Das, 2016; Roundtree et al., 2017). The levels of m6A in RNA are regulated by specific enzymes, i.e., the RNA methyltransferases and demethylases. The most widespread mRNA methyltransferase is the enzyme complex METTL3/METTL14/WTAP (methyltransferase-like 3/14 and Wilms’ tumour 1-associated protein), called also the “writer” (Liu et al., 2014; Meyer & Jaffrey, 2017). In this complex, METTL3 is the catalytic subunit. Another m6A mRNA methylation catalysing enzyme is the single subunit methyltransferase-like protein 16 (METTL16) (Brown et al., 2016; Satterwhite & Mansfield, 2022). The mRNA m6A demethylases, called “erasers”, are the fat mass and obesity-associated protein (FTO) (Jia et al., 2011) and the AlkB family member 5 (ALKBH5) (Zheng et al., 2013). The fate of the RNA in post-transcriptional processes is also governed by the so-called “reader” proteins that recognize specific m6A-modified RNA sequences. Several m6A mRNA reader proteins have been identified (Meyer & Jaffrey, 2017; Patil et al., 2018), including YTHDF1 (YTH N6- Methyladenosine RNA Binding Protein 1), YTHDF2, YTHDF3, YTHDC1 (YTH domain-containing protein 1) and YTHDC2 (Sikorski et al., 2023; Wojtas et al., 2017). These three types of proteins, i.e., writers, erasers and readers collectively coordinate the m6A mRNA methylome in the eukaryotic cell.

The m6A mRNA modification has important functions in brain development, neuronal signalling, and neurological disorders (Angelova et al., 2018; K. Du et al., 2019; Jung & Goldman, 2018; Livneh et al., 2020; Noack & Calegari, 2018; Shafik et al., 2020; Widagdo & Anggono, 2018). It has been noted that the highest levels of the m6A deposition manifest in the central nervous system, where it plays major roles in stem cell differentiation, brain development, and neurodevelopmental disorders (Livneh et al., 2020). Thus, by using the m6A immunoprecipitation of mRNAs in adult neural stem cells in combination with deep MeRIP-sequencing, m6A has been shown to be predominantly enriched in transcripts related to neurogenesis and neuronal development (J. Chen et al., 2019). Moreover, the m6A-dependent mRNA decay is critical for proper transcriptional patterning in mammalian cortical neurogenesis (Yoon et al., 2017). METTL3-mediated m6A mRNA methylation is also involved in cerebellar development by controlling the stability of transcripts related to cerebellar development and apoptosis and by regulating alternative splicing of pre-mRNAs of synapse-associated genes (C. X. Wang et al., 2018).

The mRNA m6A homeostasis in neurons is dysregulated in pathological conditions. For instance, Weng et al. uncovered an epitranscriptomic mechanism wherein axonal injury elevates m6A levels and signalling to promote mRNA translation, including regeneration-associated genes, which is essential for axonal regeneration of peripheral sensory neurons (Weng et al., 2018). As demonstrated on a limited Chinese Han population patient group, m6A-modified RNAs may play a role in conferring risk of major depressive disorder (T. Du et al., 2015). Different m6A-regulating enzymes are involved directly in the processes of learning and memory. Through its binding protein YTHDF1, m6A promotes protein translation of target transcripts in response to neuronal stimuli in the adult mouse hippocampus, thereby facilitating learning and memory (Shu et al., 2021). Similarly, the m6A mRNA demethylase FTO is expressed in adult neural stem cells and neurons and displays dynamic expression during postnatal neurodevelopment. Deficiency of FTO could reduce the proliferation and neuronal differentiation of adult neural stem cells *in vivo*, leading to impaired learning and memory (Li et al., 2017). In addition, the inactivation of the *Fto* gene in mice impairs dopamine receptor type 2 (D2R) and type 3 (D3R)-dependent control of neuronal activity and behavioural responses (Hess et al., 2013). The m6A homeostasis in neurons is controlled by more than one regulator. Thus, it was found that global m6A modification of mRNAs is down-regulated by 6- hydroxydopamine (6-OHDA) in rat pheochromocytoma PC12 cells originating from adrenal medulla and in the striatum of 6-OHDA treated rat brain that is the animal model of Parkinson’s disease (PD). The reduction of the m6A level in PC12 cells by overexpressing FTO demethylase, or by m6A inhibitor cycloleucine induced the expression of N-methyl-D-aspartate (NMDA) receptor 1, and elevated oxidative stress and Ca^2+^ influx, resulting in PC12 cell apoptosis (X. Chen et al., 2019). Recent data show that in 1-methyl-4-phenyl-1,2,3,6- tetrahydropyridine (MPTP) mouse model of PD, the mRNA m6A methyltransferase METTL3 is down-regulated in the striatum where nigrostriatal DA neurons are projecting (Z. Yu et al., 2022). In 86 individuals with PD and 86 healthy controls the levels of m6A and mRNA levels of METTL3, METTL14, and YTHDF2 in patients with PD were significantly lower compared to the healthy controls (He et al., 2023). These data indicate that dysregulation of the m6A can possibly be related to neurodegeneration in PD. However, data demonstrating that m6A system is regulating neuronal survival and neurite outgrowth and is involved in neuroprotection are still missing. Why m6A can be important for dopamine (DA) neurons that are degenerating in PD? DA neurons have special morphology with massive arborization of the dopaminergic axons. Up to 100,000 synaptic contacts per single rat midbrain DA neuron has been detected and the number may be higher for human DA neurons (Matsuda et al., 2009). Maintenance of this nigrostriatal network of DA neurons requires a lot of energy, well-functioning axonal transport, and protein synthesis. In this context it should be noted that many of the axonal proteins are not transported from the cell body to the axons but are rather synthesized on axonal ribosomes (Biever et al., 2020). This requires massive mRNA transport from the cell nucleus, where mRNAs are synthesized, to the distal axons putting DA neurons in a very special position. It should be also noted that m6A modification of mRNA may also impact organelle trafficking and neurogenesis in the axons. Taken together, therefore, it is logical to postulate that m6A mRNA methylation regulating mRNA stability, translation and transport plays an important role in the maintenance and functioning of DA neurons. This hypothesis was supported by our recent finding that demethylase FTO and ALKBH5 inhibitors supported the survival of mouse embryonic DA neurons in culture (Selberg et al., 2021).

Recently, we have demonstrated that the eukaryotic m6A mRNA methylation can be enhanced by small molecule METTL3/14 activator compounds that with high affinity and specificity stimulate the enzyme’s catalytic activity (Selberg et al., 2019). Since downregulation of m6A occurs in the PD patients, PC12 neuronal death and FTO and ALKBH5 inhibition promote cell survival we hypothesized that METTL3/14 activators may be neuroprotective and neuroregenerative for DA neurons in culture and in animal models of PD.

## Results

### METTL3/14 activators

The unique small-molecule activators of the m6A mRNA methylation have been discovered and their activity and specificity characterized (Selberg et al., 2019). The structures of four most potent compounds, C1, C2, C3 and C4 are shown in Fig. 1a. The most efficient compound, C4 has low molecular weight and very good water solubility (> 100μM) (Fig. 1b). The compounds do not inhibit hERG channel, monoamine oxidase or cytochrome P450 (Fig. 1c). The cytotoxicity screening on the number of endpoints including cell count, nuclear size, DNA structure, cell membrane permeability, mitochondrial mass, mitochondrial membrane potential (Δψm), and cytochrome c release, did not show any aberrations up to the 200 μM concentration of compounds (Fig. 1c). The compounds have good water solubility (>100μM), and the half-life of the best compound (C4) is t1/2 = 354 minutes in human plasma and 60.5 minutes in rat plasma (Fig. 1d).

**Fig. 1:**
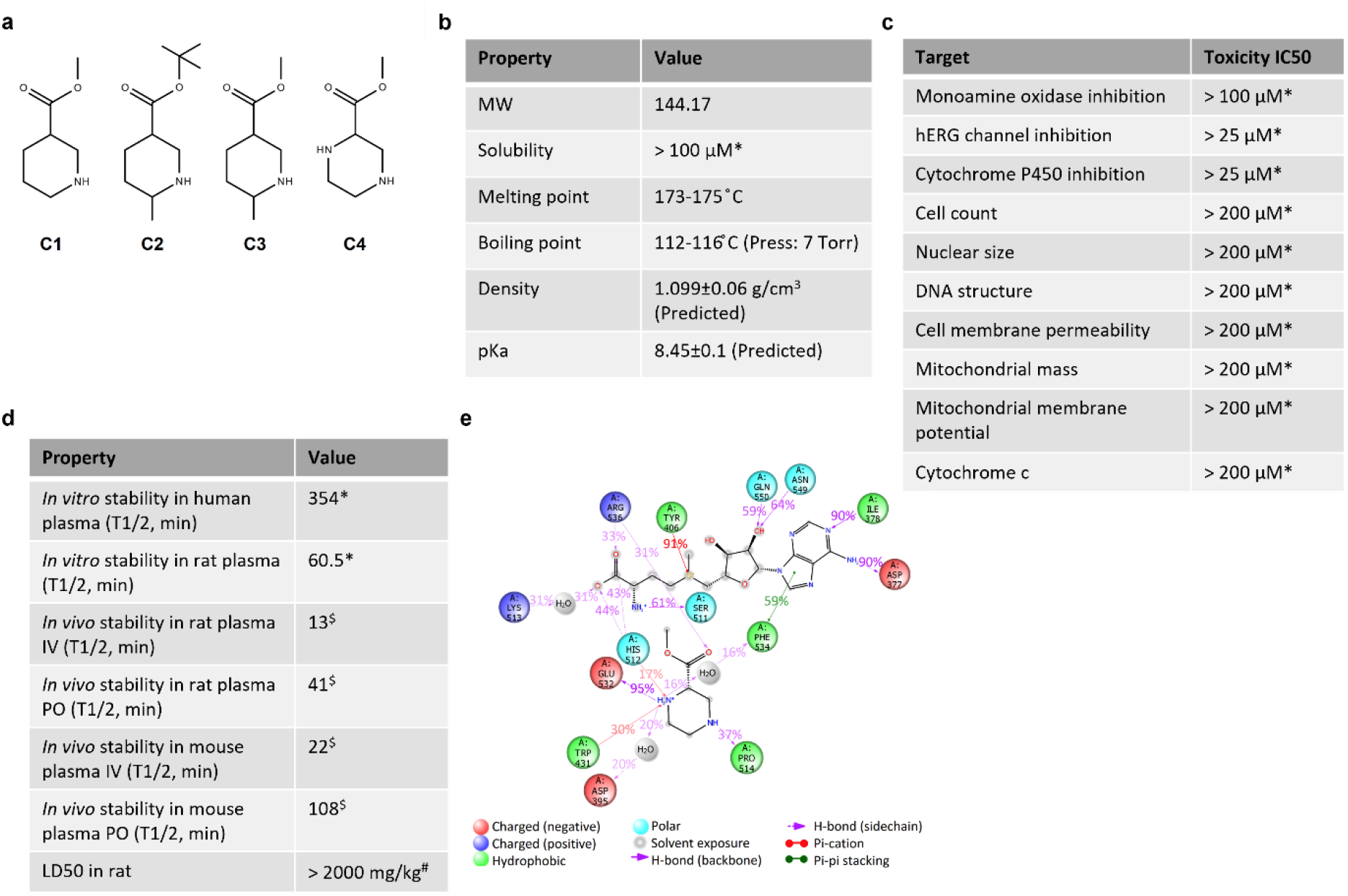
Characterisation of METTL3/14 activator C4. **a,** Chemical structure of compounds C1-C4. **b,** Properties of C4. **c,** Toxicity screen results of C4. **d,** Stability of C4. **e,** 2D summary diagram of the 50 ns molecular dynamics calculated contacts between METTL3/METTL14, C4 and SAM. Interactions that occur more than 15% of the simulation time are shown. (performed by: *Cyprotex, ^$^Admescope, ^#^Creative Bioarray)

### Modelling of the possible mechanism of action of METTL3/14 activator C4

In our previous work (Selberg et al., 2019), we hypothesized that the METTL3/14 activator C4 is binding in the close vicinity of S-adenosylmethionine (SAM) and its carboxylate group is co-ordinated with the positively charged S+ atom of the SAM. Thus, potentially increasing the binding energy of the SAM due to the electrostatic charge-dipole interaction as well as reducing the methylation reaction activation barrier (Hussain et al., 2015). To further evaluate the possible mechanism of action, we carried out the molecular dynamics (MD) study of the interactions between the catalytic domain of the METTL3/METTL14 (PDB ID: 5K7W) (P. Wang et al., 2016) and SAM (PDB ID: 5IL1) (X. Wang et al., 2016) both in the presence and absence of the METTL3/14 activator C4. The analysis of the MD simulations results show that compound C4 has a great impact on the binding mode of the SAM in the catalytic domain of METTL3/METTL14 (Fig. 1e; Suppl. Fig. S1a). Namely, in the presence of compound C4, the contacts with Phe534 and Asn549 amino acid residues that allow to hold the active position of the adenine ring (P. Wang et al., 2016) are changed. The duration of p-p stacking between the sidechain of Phe534 is slightly decreased, from 75% to 59% of simulation time. However, the sidechain of Asn549 has a strong hydrogen bond with hydroxyl group on the adenosine ribose in the presence of C4 (lasting 64% of the simulation time), instead of the several hydrogen bonds and water bridges with backbone. The contact with the conserved Asp395 of the ^395^DPPW^398^ catalytic motif is also changed upon binding with the activator C4. Instead of the long-lasting hydrogen bond between the hydroxyl group on the ribose of the SAM and the sidechain of Asp395, a short-lasting water bridge appears between the N atom of the piperazine ring of the ligand (Fig. 1e; Suppl. Fig. S1a). In addition, the nitrogen atom of the C4 piperazine ring takes over the place of the positively charged SAM amino group in a long-lasting hydrogen bond with the sidechain of Glu532 amino acid residue whose importance in the SAM coordination has been confirmed by mutational analysis (X. Wang et al., 2016). The most significant change in the binding mode of SAM with the METTL3/METTL14 catalytic domain in the presence of C4 is appearing in the long-term specific p-cation interaction between the S+ atom of SAM and an evolutionarily conserved Tyr406. In the absence of C4, this specific contact is extremely short (5% of the simulation time) (Suppl. Fig. S1b), while in the presence of the activator, this contact persists throughout the entire MD simulation (91% of the simulation time) (Fig.1e). The stability of the MD simulation and results obtained were confirmed by additional MD simulation with a length of 25 ns with the same complexes as well as using another crystal structure of the METTL3/METTL14 complex (data shown at request). Thus, based on the MD simulation results, upon binding with METTL3/METTL14/SAM complex, activator C4 takes over and boosts some of the specific interactions between SAM and METTL3/METTL14 thus promoting more efficient transfer of the methyl group to the adenosine in N6-position.

### METTL3/14 activators support the survival and promote neurite outgrowth of dopamine neurons

Two main reasons prompted us to test the effect of METTL3/14 activators on DA neurons. Firstly, it has been earlier shown that m6A levels are decreased in animal models of PD (X. Chen et al., 2019; Z. Yu et al., 2022) and in the sera of PD patients (He et al., 2023). Secondly, midbrain DA neurons have massive neurite branching (Matsuda et al., 2009) predicting the importance of mRNA stability and transport, and consequently the significance of m6A for the maintenance and functioning of these neurons. To test the effect of METTL3/14 activators on the survival and neuroprotection of mouse embryonic DA neurons, we first studied and verified the mRNA expression of *METTL3/14* and several other members of the m6A system in mouse DA neurons using RT-qPCR (Suppl. Fig. S2a) and immunocytochemistry (ICC). METTL3 and m6A are present in neurons as demonstrated by immunostaining of human induced pluripotent stem cell-derived (iPSC) DA neurons and embryonic day E13 mouse midbrain DA neurons with m6A and METTL3 antibodies (Fig. 2a). It is important to note that both METTL3 and m6A immunoreactivity is mostly detectable in DA neuron cell bodies, but also in axons (Fig. 2a). Based on earlier published data (J. Chen et al., 2019; Z. Yu et al., 2022) and our own data on FTO and ALKBH5 inhibitors (Selberg et al., 2021), we hypothesized that the activation of mRNA m6A methyltransferase METTL3/14 in DA neurons could counteract their apoptotic cell death and support the survival of neurons. To test this hypothesis, we assessed the survival-promoting effect of METTL3/14 activators C1-C4 on the mouse midbrain DA neurons after inducing their apoptosis by growth factor deprivation (Kovaleva et al., 2023; L. Y. Yu et al., 2008). Glial cell line-derived neurotrophic factor (GDNF) has been shown to protect and rescue cultured embryonic DA neurons from growth factor deprivation-induced apoptosis, as well as 6-OHDA-induced cell death *in vitro* and *in vivo* (Eesmaa et al., 2021; Kovaleva et al., 2023; Lindholm et al., 2007; L. Y. Yu et al., 2008). We, therefore, assessed the neuroprotective ability of different concentrations of selected METTL3/14 activators C1-C4 in cultured growth factor deprived DA neurons. Human recombinant GDNF (100 ng/ml) or a condition without any neurotrophic compound added were used as positive and negative controls, respectively. Growth factor deprivation caused cell death by approximately 40%. The results, expressed as percentage of cell survival compared to GDNF-maintained neurons for the METTL3/14 activators, show that C3 and C4 (Fig. 2b) more efficiently than C1 and C2 (Suppl. Fig. S2b) rescue mouse DA neurons. METTL3/14 activators C3 and C4, similarly to GDNF, dose-dependently protected mouse DA neurons in culture from growth factor deprivation-induced cell death already at 10 nM, and significant protection was observed also at 100 nM and 1 μM concentrations, and even at 1 μM concentration no signs of toxicity of the tested compounds were observed (Fig. 2b). The neuroprotective ability of C3 and C4 were then tested in neurotoxin 6-OHDA-treated mouse DA neurons. Again, C3 and C4 protected DA neurons from 6-OHDA-induced cell death at two C3 and C4 concentrations tested – at 100 nM and 1 μM concentrations (Fig 2c). These data demonstrate that METTL3/14 activators can protect and rescue mouse DA neurons in two different cell death paradigms, indicating that mRNA m6A plays crucial role in neuronal survival and maintenance.

**Fig. 2:**
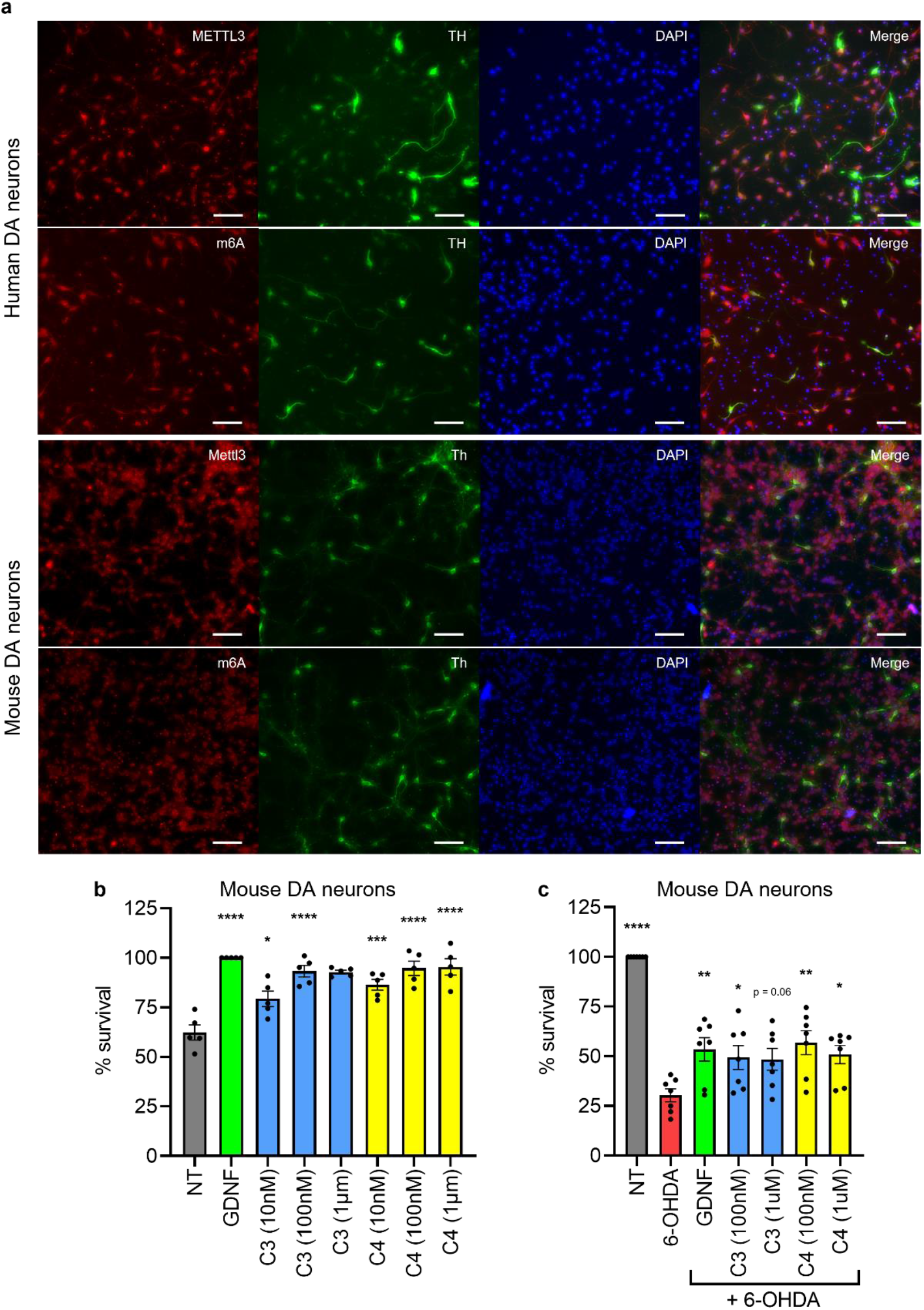
Effect of METTL3/14 activation on mouse DA neurons. **a,** Representative images of human and mouse DA neurons stained with METTL3 (red), m6A (red), TH (green) antibodies and counterstained with DAPI (blue), scale bar 100µm. **b,** Survival of mouse DA neurons in growth factor deprivation assay with C3 or C4 compared to GDNF (100ng/ml). **c,** Survival of mouse DA neurons with GDNF (100ng/ml) C3 or C4 in the presence of 6-OHDA (15µM). *p<0.05, **p<0.01, ***p<0,001, ****p<0.0001

Our earlier data demonstrated that the demethylase FTO inhibitor F-1-7 (compound 2) and demethylase ALKBH5 inhibitor AL-1-15 (compound 8) protected mouse DA neurons at 10 nM concentrations (Selberg et al., 2021). Since RNA demethylase inhibitors and methyltransferase activators increase or maintain mRNA m6A levels i.e., work in the same direction, we asked whether these compounds have addictive effects. We tested the neuroprotective effects of FTO inhibitor F-1-7 and demethylase ALKBH5 inhibitor AL-1-15 in combination with METTL3/14 activator C4 on 6-OHDA treated mouse DA neurons. Results presented on Suppl. Fig. S2c show that m6A methyltransferase activators and demethylase inhibitors when tested together at 100 nM or 1 µM concentrations show no additive effects.

Direct survival-or neurite outgrowth-promoting effects of GDNF on human DA neurons have not been described. The role of mRNA m6A on the survival and neurite outgrowth of human DA neurons has not been studied. As an important step towards clinical translation, we tested the survival promoting and neurite outgrowth inducing effects of GDNF and METTL3/14 activator C4 on human DA neurons. We have used human iPSC-derived DA neurons from FUJIFILM Cellular Dynamics Inc. (see Methods section) and worked out conditions to grow and assay these neurons in the medium lacking growth factors and proteins. We have established a survival assay with 6-OHDA treatment (Fig. 3a) and the neurite outgrowth assay. Results shown on Fig. 3a clearly show that METTL3/14 activator C4 dose-dependently protects human DA neurons from 6-OHDA-induced neuronal cell death. The neuroprotective effect of GDNF is at 100 ng/ml (∼3 nM) and C4 compound protects neurons already at 10 nM. At the PD diagnosis a significant number of nigrostriatal DA neurons have degenerated axons. Thus, in addition to the survival-promoting and neuroprotective activity, another important property of an efficient PD drug is the ability to regenerate the axons and dendrites of DA neurons. For these reasons we tested the capacity of GDNF and METTL3/14 activator C4 to induce neurite outgrowth from cultured human DA neurons. Results presented in Fig. 3b-d clearly show that GDNF significantly induced outgrowth of neurites and increased the number of processes per cell in human DA neurons. What is more, C4 shows spectacular neurite outgrowth and increase in number of processes (Fig. 3b-c). Although there was no statistically significant change in the number of neurite branches, a positive trend was observed with both GDNF and C4 (Fig. 3d). Taken together, this is the first demonstration that GDNF can protect human DA neurons against cell death, also inducing axonal growth. Most importantly, these data for the first time show that METTL3/14 activators have neuroprotective and neurite outgrowth-promoting activity in human neurons. Since METTL3/14 activators protect DA neurons from cell death it is logical to predict that METTL3/14 inhibitors have an opposite effect. We used a potent METTL3/14 inhibitor STM2457 that specifically binds to the enzyme and inhibits its catalytic activity with IC_50_ at 16.9 nM concentration. At 2-5 µM concentration STM2457 also inhibits METTL3/14 catalytic activity in cells (Yankova et al., 2021). The binding of STM2457 to the METTL3/14 complex was confirmed with a K_d_ of 754 nM in microscale thermophoresis assay (Fig. 3e). When added to the media of mouse DA neurons, STM2457 dose-dependently kills these neurons with IC_50_ around 25μM (Fig. 3f). These data demonstrate that METTL3/14 activators promote DA neurons’ cell survival, whereas inhibitor of that enzyme triggers neuronal cell death emphasising the importance of m6A in the life and death of DA neurons.

**Fig. 3:**
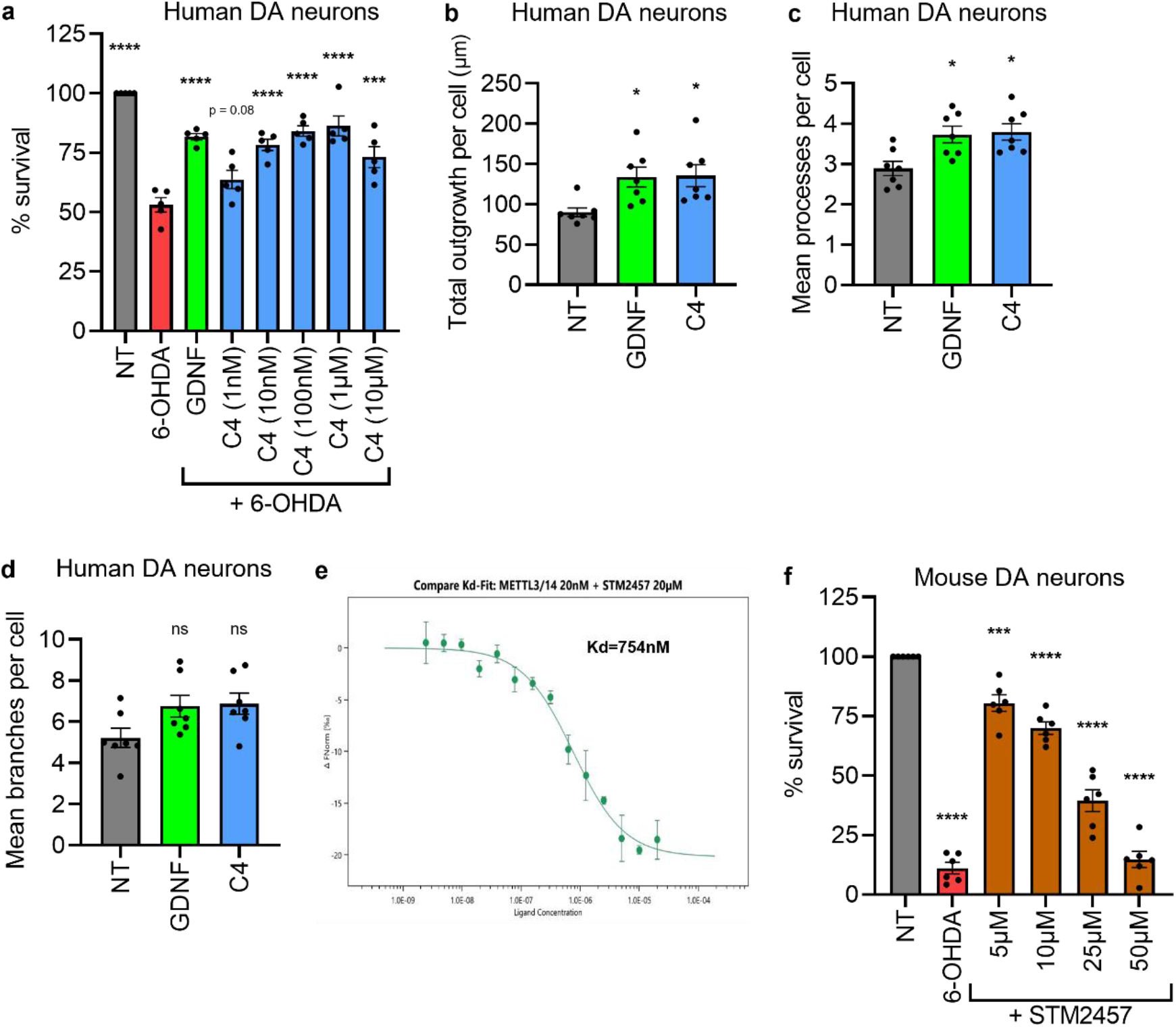
Effect of METTL3/14 activation on human DA neurons. **a,** Survival of human DA neurons with GDNF (100ng/ml) or C4 in the presence of 6-OHDA (50µM). **b-d,** Human DA neuron neurite outgrowth assay demonstrating (**b**) total outgrowth in µm, (**c**) mean processes and (**d**) mean branches per cell following GDNF (100ng/ml) or C4 (1µM) treatment. **e**, Microscale thermophoresis assay showing binding of STM2457 with METTL3/14 complex **f,** Survival of mouse DA neurons following treatment with 6-OHDA (15µM) or STM2457 compared to NT (nontreated). Data are presented as mean±SEM. *p<0.05, **p<0.01, ***p<0,001, ****p<0.0001

After demonstrating anti-apoptotic and neuroprotective effects of METTL3/14 activator on mouse and human DA neurons, we asked whether these compounds are specific for DA neurons or they promote the survival of other types of neurons as well. This is important because in systemic delivery these compounds can influence in addition to midbrain neurons also other brain neurons, and especially peripheral neurons. We tested the effects of C4 on mouse postnatal superior cervical ganglion (SCG) sympathetic neurons and embryonic dorsal root ganglion (DRG) sensory neurons. Neuronal cell death was induced by treatment with endoplasmic reticulum (ER) stress-inducing agent tunicamycin (2 µM). Compound C4 was added at 100 nM and 1 μM concentrations. Unexpectedly METTL3/14 activator C4 had no survival-promoting or neuroprotective effects on SCG and DRG neurons (Suppl. Fig 2d). Similar results were obtained when SCG and DRG neuronal cell death was triggered by growth factor deprivation (data not shown). We hypothesized that either the C4 target METTL3/14 is not expressed in SCG sympathetic and DRG sensory neurons or C4 is not penetrating the cell membrane of these neurons. To distinguish between these two possibilities, we first analysed the presence of METTL3 and m6A in these neurons using ICC. Our results clearly show that both SCG and DRG neurons prominently express METTL3 and interestingly its expression is detectable also in neuronal axons (Suppl. Fig. S2e). When we added C4 to the media of SCG or DRG neurons we did not observe survival-promoting or neuroprotective effects even at 1 μM concentration. To test whether C4 lacks the effects on these neurons because it is unable to penetrate the cell membrane of these neurons, we took advantage of the neuronal microinjection technique. Microinjection of the mRNA m6A methyltransferase METTL3/14 activator C4 can rescue tunicamycin-treated ER-stressed SCG and DRG neurons from death (Suppl. Fig. S2f). C4 was as efficient as mesencephalic astrocyte-derived neurotrophic factor (MANF) that was used as the positive control (Eesmaa et al., 2021; Kovaleva et al., 2023). These results clearly demonstrate that the METTL3/14 activator C4 is not specific for DA neurons, but it does not affect DRG sensory and SCG sympathetic neurons when added to the culture media even at as high concentration as 1 μM. Our data indicate that due to the differences in the structure of the plasma membrane, C4 simply is unable to enter SCG and DRG neurons in culture.

### Analysis of mRNA expression in dopamine neurons after treatment with METTL3/14 activator C4

We analysed cultured mouse dopamine neurons treated with 6-OHDA, with METTL3/14 activator C4 alone and with 6-OHDA plus C4 compound. Untreated DA neurons served as a control. We carried out the mRNA expression analysis of these neurons using qPCR for the selected genes belonging to the following groups: growth factors, growth factor receptors, key enzymes in m6A mRNA methylation, and key enzymes of dopamine synthesis and metabolism. We anticipated to see some changes in the expression of these genes after 6-OHDA and C4 treatment. We isolated RNA from our DA neuron cultures that contain in addition to the DA neurons also striatal neurons that are innervated by DA neurons and glial cells. We hoped to monitor this way also changes in the levels of mRNAs encoding for growth factors important for the survival and regeneration of DA neurons that are usually expressed and secreted by target neurons. We also isolated RNA from purified DA neurons. To that end we used transgenic mice that express green fluorescent protein (GFP) under the tyrosine hydroxylase (TH) promoter (Matsushita et al., 2002). DA neurons from these mice were cultured and treated as described above and then purified using fluorescent cell sorting. We estimate that the purity of DA neurons in these preparations is over 95%. qPCR analysis of the mRNAs from these neurons allows to detect mRNAs solely expressed by DA neurons. Analysis of the qPCR results clearly showed that 6-OHDA upregulates several genes (Suppl. Fig. S2a). We registered a strong upregulation of the genes encoding for the proteins known to support the survival and protect rodent and non-human primate DA neurons. Among these are the mRNAs encoding for GDF15, CDNF and MANF growth factors. From mRNAs encoding for neurotrophic factor receptors, we observed a clear trend for upregulation of GDNF binding receptor GDNF family receptor alpha 1 (GFRα1). In analysis of the mRNA levels encoding proteins that regulate dopamine synthesis, turnover and metabolism we detected clear downregulation of the DA transporter Slc6a3. Somewhat unexpectedly we detected a trend of upregulation of the dopamine receptor D1 and not D2 that was regulated by demethylase FTO (Hess et al., 2013). When we analysed the effects of C4 compound, the only statistically significant change was observed with Ret mRNA when compared to the changes in the mRNA levels induced by 6- OHDA (Suppl. Fig. S2a).

### METTL3/14 activator C4 effects in rat 6-OHDA model of Parkinson’s disease

The efficacy of METTL3/14 activator compound C4 was tested at two different doses in rat unilateral 6-OHDA model of PD (Lindholm et al., 2007; Voutilainen et al., 2009). Since we wanted to compare its efficacy directly with the “gold standard” GDNF, we used intracranial delivery of drugs. Rats were first intrastriatally injected with 6 μg of 6-OHDA (injection to three sites, 2 μg each) and two weeks later the delivery of the drugs started. We compared the effects of C4 at two doses (0.3 μg/24 h; 1.5 μg/24 h) with GDNF (3 μg/24 h) in a unilateral 6- OHDA PD model in rats using striatal delivery with Alzet osmotic minipumps. Vehicle, C4 at two doses, and GDNF were delivered for four weeks (Fig. 4a). To assess, whether C4 can restore motor impairment, we monitored amphetamine-induced ipsilateral rotations. The average number of net ipsilateral turns was in the same range in all treatment groups at 2 weeks post lesion (around 650 turns/120 min). At 6 weeks post lesion, the delivery of drugs was finished and the number of net ipsilateral turns recorded. C4 treatment at the lower dose (0.3 μg/24 h) resulted in a significant decrease in turns when compared to vehicle (Fig. 4b). C4 at the higher dose (1.5 μg/24 h) showed a positive trend (p = 0.07). Notably, GDNF did not have an effect at this time point. At week 9, C4 at the higher dose showed a significant reduction in turns when compared to vehicle, whereas GDNF and C4 at the lower dose showed a trend in reducing the ipsilateral turns. At week 12, both C4 doses exhibited a clear trend in reducing the turns (p = 0.07). GDNF, however, remained comparable to vehicle. These data clearly demonstrate that C4 efficiently and more rapidly than GDNF improved the motor behaviour of 6-OHDA lesioned rats.

**Fig. 4:**
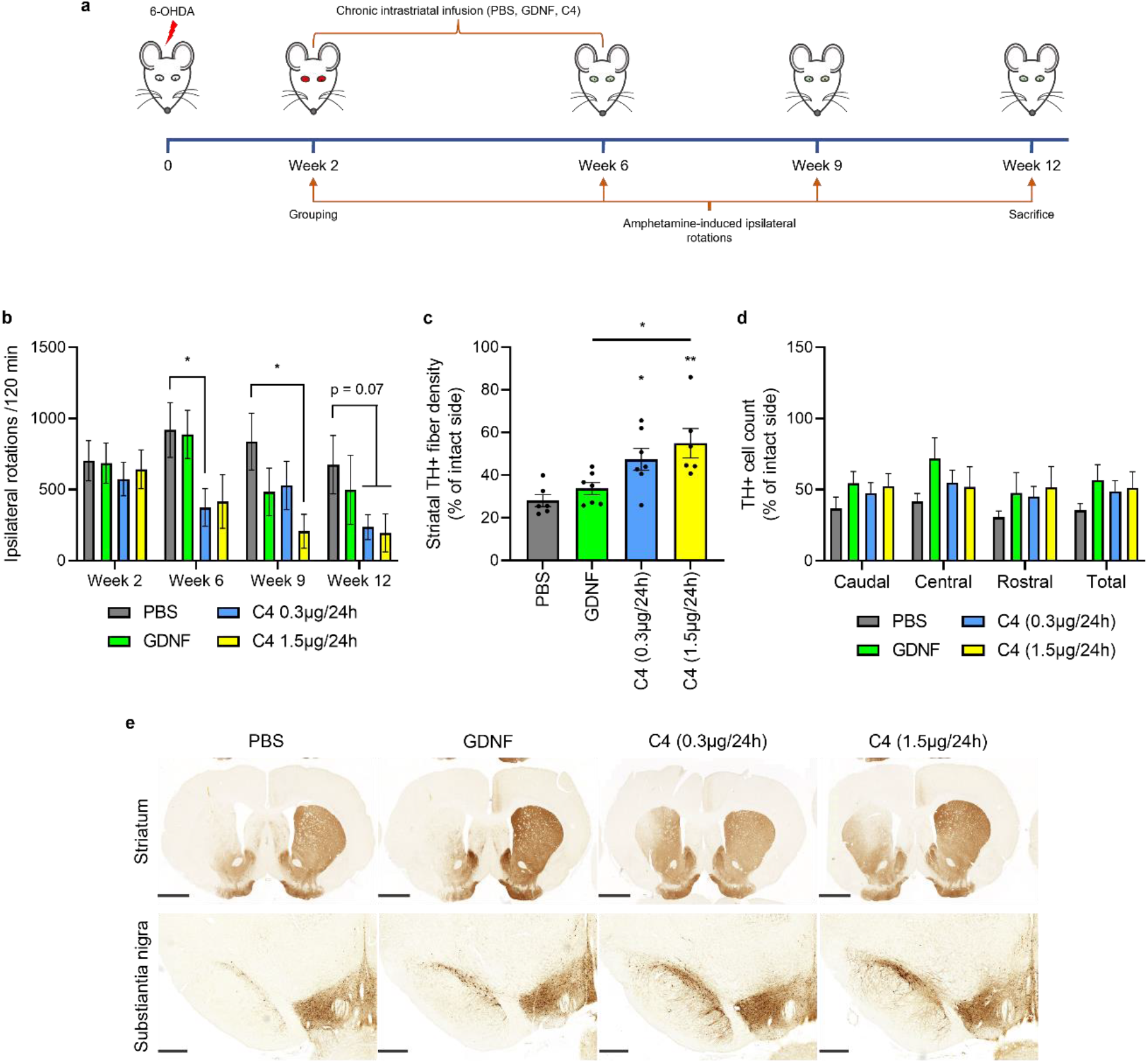
Effect of METTL3/14 activator C4 in rat 6-OHDA model of Parkinson’s disease. **a,** *In vivo* rat 6-OHDA experiment scheme. **b,** Amphetamine-induced ipsilateral rotations in rats following intrastriatal lesion with 6-OHDA (3 x 2 µg) and chronic intrastriatal infusion with GDNF (3µg/24h) or C4 (0.3µg/24h or 1.5µg/24h). **c,** TH+ fiber density of rat striata compared to the intact side. **d,** TH+ cell count in different SNpc regions. **e,** Representative TH+ fiber staining pictures, scale bars: Striatum 2mm, Substantia nigra 0.5mm. *p<0.05, **p<0.01

To evaluate the integrity of the nigrostriatal dopamine system in this neurorestoration study, coronal brain sections from the SNpc and striatum were analysed histologically after the last behavioural test i.e., 12 weeks after 6-OHDA lesion. The single unilateral injection of 6-OHDA into three sites of the dorsal striatum (3 x 2 μg) resulted in a relatively severe degeneration and loss of DA neurons: in rats treated with vehicle, the density of TH-positive fibres in the striatum was 28% in comparison to the intact side (Fig. 4c,e). Because the lesion was severe, GDNF infusion showed only a trend to increase the density of TH-positive fibres in the lesioned striatum (∼34% in TH-immunoreactivity) as compared to PBS (Fig. 4c,e). Both tested doses of C4 compound 0.3 μg/24 h and especially the dose 1.5 μg/24 h, produced a robust and statistically significant increase in the density of TH-positive fibres (47% and 55% in TH-immunoreactivity, respectively). A positive trend of GDNF and C4 on the number of TH-positive cell bodies was observed in different SNpc regions, although this remained statistically insignificant (Fig. 4d). These data demonstrate that METTL3/14 activator C4 protects and restores the function of DA neurons *in vivo* in rat 6-OHDA model of PD. Compared to GDNF, the results show that METTL3/14 activator C4 acts faster and is more potent in protecting and regenerating the axons of DA neurons. Effects of C4 and GDNF on DA neuron cell bodies are similar.

## Discussion

### METTL3/14 activator enhances the interaction between METTL3/14 and RNA substrate

In 2019, we reported the discovery of unique activators of the METTL3/METTL14/WTAP complex using computer-aided drug design (CADD) (Selberg et al., 2019). It was experimentally shown that these compounds act as artificial co-factors to the METTL3/14 complex (METTL3/14 activators) and enhance the binding efficiency of the methyl group donor SAM by several orders of magnitude. In this work, we report the results of the in-depth modelling of the possible mechanism of action of METTL3/14 activator C4 at molecular level using MD simulations, one of the most powerful CADD tools (Lin, 2022). This method makes it possible not only to consider the flexibility of the receptor and ligand that is crucial for the correct prediction of drug binding but also to evaluate the protein-ligand contacts in real time (De Vivo et al., 2016). According to the present results, the most significant change upon C4 binding is related to the strengthening of the p-cation contact between Tyr406 of METTL3/14 and the S+ atom of SAM. The presence of this contact promotes the proper interaction of METTL3 with the RNA substrate near the adenine to be methylated and catalyses the methyl transfer (P. Wang et al., 2016). This finding is in line with our previous experimental data and it can be concluded that at the molecular level the mechanism of action of compound C4 is related to boosting the methyl group transfer process by increasing the strength of binding between the METTL3/14 catalytic domain and SAM.

### Effect of C4 on mouse dopamine neurons

The mRNA m6A levels are decreased in the brain of 6-OHDA treated rodents that is the widely used animal model of PD (X. Chen et al., 2019). Decrease in the level of m6A mRNA methylation was also reported in PD patients when compared to the healthy controls (He et al., 2023). Analysis of the m6A demethylase FTO knockout mice demonstrate that FTO regulates midbrain dopaminergic signalling and affects the functioning of dopamine receptor type 2 and type 3 (Hess et al., 2013; Ruud et al., 2019). However, the role of m6A methylation on neuronal survival and maintenance, as well as neurite outgrowth and regeneration have not been described. Furthermore, direct role of m6A methylation on neuroprotection in animal models of PD has not been reported. We first found that the most potent METTL3/14 activators C3 and C4 rescue mouse DA neurons already at 10 nM concentration when added to the media in growth factor deprivation cell death paradigm. The efficacy of the compounds was comparable to “gold standard” GDNF. These results clearly demonstrate that m6A methylation activators protect DA neurons from the type of neuronal cell death that also occurs in PD patients. Namely reduced levels of GDNF, that was deprived from DA neurons in these experiments, have been reported in PD patent brains (Virachit et al., 2019). We then tested the neuroprotective capacity of m6A activators in mouse neurons treated with neurotoxin 6-OHDA. This is the widely used *in vitro* model of PD, where neurons die because of the 6-OHDA-induced oxidative and ER stress. C3 and C4 demonstrated strong neuroprotective ability against 6-OHDA induced neuronal cell death. In these experiments the efficacy of C4 was somewhat more pronounced than that of C3. Collectively these data demonstrate that m6A activators support the survival and protect neurons against 6-OHDA-induced neurodegeneration.

### METTL3 inhibitor induces the death of DA neurons in culture

Novel effective inhibitors binding to and blocking the catalytic activity of METTL3/14 have been recently developed (Yankova et al., 2021). Authors showed that the compound STM2457 is a potent inhibitor of METTL3/14 catalytic activity with an IC_50_ of 16.9 nM and it significantly inhibited METTL3/14 activity in several cell lines already at micromolar concentrations. In a series of eloquent *in vivo* experiments, the authors also demonstrate that inhibition of METTL3 is a potential therapeutic strategy against acute myeloid leukaemia and can be an exciting new avenue to develop anticancer therapy. We anticipated that while METTL3/14 activators protect and maintain DA neurons, inhibition of m6A methylation may trigger cell degeneration and death. We first confirmed that STM2457 preparation used in our microscale thermophoresis (MST) experiments binds with high affinity to METTL3/14 complex. K_d_ determined in our experiments is higher than that found by SPR in original publication describing METTL3/14 inhibitor STM2457 (Fig. 3e) (Yankova et al., 2021). We have observed significantly higher K_d_ values in MST also for GDNF and its receptor’s GFRα1 interaction. We then tested the effect of STM2457 on the survival of cultured mouse DA neurons. Indeed, STM2457 in a dose-dependent manner triggered the death of DA neurons further supporting the point of view that m6A methylation of mRNA plays a crucial role in neuronal survival.

### METTL3/14 activator C4 effects on the survival and neurite outgrowth of human dopamine neurons

There is increasing evidence that rodent DA neurons differ from human DA neurons both in morphology, but also in molecular characteristics. iPS cell technology has enabled to develop human DA neurons and test them in tissue culture and *in vivo*. Several teams are currently testing stem cell-derived human DA neurons in clinical trials for PD and we are looking forward to hearing soon the results. Despite of multiple activities on human DA neurons, the effects of GDNF and other neurotrophic factors in 6-OHDA models of PD have not been tested in human DA neurons. What is more, also GDNF effects on the ability to induce neurite outgrowth and axonal regeneration have not been tested in human DA neurons. It is obvious that m6A methylation effects on human DA neurons have not been tested at all. Using iPS cell-derived human DA neurons, we first established the conditions to test growth factor effects in 6-OHDA treated neurons. Our results show that GDNF dose-dependently protects human DA neurons in culture against neurotoxin 6-OHDA induced cell death. What is even more interesting is that C4 already at 10 nM protected human DA neurons showing for the first time that m6A mRNA methylation is crucial also for human DA neuron survival. As mentioned above, GDNF effects on neurite outgrowth of human DA neurons have been never studied. We first worked out the conditions to monitor growth factor-induced neurite outgrowth from human DA neurons. We used axonal markers and monitored the axonal branching as well as cumulative neurite outgrowth, because both parameters are important for neuroregeneration of DA neurons in PD patients. We observed that as in rodent DA neurons *in vitro* and in *in vivo*, GDNF can induce neurite outgrowth and branching in human DA neurons (Fig. 3b-d). METTL3/14 activator C4 induced a prominent neurite outgrowth and increased the number of processes of human DA neurons being at least as effective as GDNF. These results demonstrate that in addition to the neuroprotection, m6A activator C4 has also neuroregenerative activity by inducing the neurite outgrowth and increasing the number of processes of human DA neurons. These results imply that m6A activators have a significant role on axonal regeneration and the compounds fulfil two important criteria for the drugs of disease-modifying treatment of PD – they protect and regenerate DA neurons.

### C4 effects of peripheral sympathetic and sensory neurons

Whether m6A activators are specific to DA neurons, is one of the most important questions. To test that, we treated early postnatal sympathetic SCG neurons and late embryonic sensory DRG neurons with tunicamycin to trigger cell death and then with C4 compound. Surprisingly m6A activator compounds C4 had no survival-promoting effects on these neurons. At the same time, MANF (and also nerve growth factor, NGF; data not shown) efficiently rescued the sensory and sympathetic neurons in these experiments. C4 was tested in a wide range of concentrations that were several orders of magnitude above the EC_50_ of the compounds (Selberg et al., 2019). These results raised two possible scenarios for the explanation. Either SCG and DRG neurons do not express the target proteins METTL3/14 or the C4 is unable to penetrate the cell membrane of these neurons. We tested both possibilities and found that METTL3/14 is expressed at relatively high levels in both types of peripheral neurons. Furthermore, when we then used neuronal microinjection technique and microinjected C4 into SCG and DRG neurons, we found that C4 compound protected these neurons from tunicamycin induced cell death. These results clearly demonstrate that C4 can protect and rescue both SCG and DRG neurons but is unable at even high concentrations of C4 to pass thorough the plasma membrane of these neurons. These results suggest that C4 is not specific for DA neurons, but at least peripheral sensory and sympathetic neurons have a cell membrane that does not allow the C4 to penetrate these cells. The *in vivo* implications of these results are rather clear. We anticipate that m6A activator C4, when delivered systemically, would not affect peripheral sensory and sympathetic neurons.

### C4 protects and repairs dopamine neurons in rat 6-OHDA model of Parkinson’s disease

Neurotoxin 6-OHDA-induced degeneration is one of the most commonly used PD models. If the potential drug candidate is injected several weeks after 6-OHDA then the model quite well recapitulates the human disease situation. GDNF has been widely used to test its ability to protect and repair DA neurons in animal models of PD. In moderate neurotoxin model, i.e., when 30-70% of the DA neurons are still alive after lesion, GDNF protects and repairs DA neurons in the 6-OHDA animal model of PD. However, in clinical trials when more than 80% of the human DA neurons are lost, the neuroprotective effects of GDNF are modest. We used the rat 6-OHDA model of PD and first injected 6-OHDA, and two weeks later, when about 50 % of the DA neurons have been lost, like the situation of PD patients at the moment of PD diagnosis, we started to infuse m6A activators. We used intracranial chronic delivery mostly because we wanted to compare the effects of C4 with GDNF that works well in moderate neurotoxin animal models and has shown promise also in clinical trials (Whone et al., 2019). We used Alzet minipumps connected with plastic tubing to brain cannulas to deliver vehicle, GDNF as positive control, and two doses of m6A activator C4 into the striatum to test their neurorestorative activity. Since C4 infusion was started 2 weeks after 6-OHDA treatment when the rats had clear motor symptoms and DA neurons’ neurodegeneration started, we can conclude that C4 has neurorestorative activity. Importantly, C4 at both lower and higher doses significantly restored locomotor activity at 6 and 9 weeks after 6-OHDA lesion, respectively, whereas GDNF showed a positive trend at week 9. At week 12, C4 at both doses showed a strong trend in reducing amphetamine induced ipsilateral rotations of rats demonstrating that C4 acts faster and more efficiently than GDNF and its effect is long lasting i.e., can be detected even 6 weeks after the drug delivery was stopped. Postmortem analysis of TH-positive striatal fibres demonstrates that only 28% of the fibres can be detected 12 weeks after the 6-OHDA lesion. In this severe 6-OHDA lesion paradigm GDNF has no statistically significant effect, but C4 at both does has a robust effect. C4 efficiently protected TH-positive fibres, but it is still unclear whether it can also regenerate DA neuron axons *in vivo*. This is very likely because C4 induced neurite outgrowth of cultured DA neurons. C4 at both doses and GDNF showed a positive trend in protecting DA neuron cell bodies from 6-OHDA induced cell death, however due to high variability this remained statistically insignificant. Here the effects of GDNF and METTL3/14 activator C4 are similar suggesting that C4 impacts more on axonal growth and regeneration than to the neuroprotection of the cell bodies. Further studies are needed to understand this difference in GDNF and C4 effects.

### qPCR analysis of genes in dopamine neurons – possible implications to the mode of action of m6A activator compounds

Midbrain DA neurons form a huge number of synaptic contacts with target cells. Since many proteins are synthesized in the axons and dendrites this requires rather massive RNA transport form the nucleus, where RNA is synthesized, to various locations in the axons and dendrites. We hypothesized that m6A methylation plays a crucial role in the regulation of mRNA stability and transport to maintain and regulate large number of DA neurons’ synapses. Predictably the levels of some of the mRNAs should be increased and vice versa some of the mRNAs, perhaps those encoding axonal growth inhibiting or apoptosis inducing proteins, will be downregulated? Our qPCR analysis of four groups of transcripts, encoding for neurotrophic growth factors, growth factor receptors, m6A regulating enzymes and proteins, and enzymes regulating dopamine synthesis and metabolism, demonstrated strong upregulation of many mRNAs after 6-OHDA treatment. Somewhat unexpectedly, no clear changes in the levels of selected mRNAs were found in 6-OHDA-treated cultured DA neurons after C4 addition. Nevertheless, it is reasonable to argue that mRNAs encoding proteins that impact axonal growth and regulation might be influenced by C4, as this is also supported by our *in vitro* and especially by *in vivo* data demonstrating that C4 is a strong inducer of DA neuron’s axonal growth.

## Conclusions

Several important conclusions can be drawn from our results. Firstly, this is the first demonstration that m6A mRNA methylation supports the survival and regeneration of the axons of mouse and human DA neurons, and perhaps CNS neurons in general. Secondly, this m6A mRNA methylation is a completely novel pathway regulating neuronal survival and regeneration that opens an entirely new avenue for drug development. Thirdly, tested METTL3/14 activators showed no toxicity on cells, neurons and in animal experiments suggesting that these compounds can be delivered systemically. Fourthly, having worked for more than 20 years on DA neurons we have not seen earlier a small molecule that supports the survival and protects DA neurons more efficiently than GDNF. Although C3 and C4 already meet the main criteria of lead compounds from the medicinal chemistry point, we see clear possibilities to optimize these compounds.

## Methods

### Molecular dynamics

The crystal structure of the catalytic domain of METTL3/METTL14 complex with S-adenosylhomocysteine (PDB ID: 5K7W) (P. Wang et al., 2016) and the crystal structure of SAM-bound METTL3/METTL14 complex (PDB ID: 5IL1) (X. Wang et al., 2016) were used for the MD simulations. The binding mode of the METTL3/14 activator C4 was predicted by molecular docking using AutoDock Vina 1.1.2 (Trott & Olson, 2009). The ligand conformation having the strongest docking energy was used in the further study. The MD simulations were carried out using Desmond simulation package of Schrödinger LLC (Bowers et al., 2006; D. E. Shaw Research, New York, 2020; Schrödinger, 2021). For each studied complex, the simulation lengths were 50 nanoseconds (ns) with a relaxation time of 1 picosecond (ps). The simulations were carried out in cubic simple point-charge (Zielkiewicz, 2005) water box using OPLS_2005 force field parameters (Banks et al., 2005). Sodium and chloride ions were placed in the solvent to a concentration 0.15 M, and then, to achieve electro neutrality, additional ions were added to the system. The NPT ensemble with the temperature 300 K and pressure 1 bar was applied in all runs. The Martyna-Tuckerman-Klein chain coupling scheme (Martyna et al., 1992) with a coupling constant of 2.0 ps was used for the pressure control, and the Nosé-Hoover chain coupling scheme (Martyna et al., 1992) was used for the temperature control. The long-range electrostatic interactions were calculated using the Particle Mesh Ewald method (Toukmaji & Board, 1996). The cut-off radius in Coloumb´ interactions was 9.0 Å. Nonbonded forces were calculated using a reversible reference system propagation algorithm integrator where the short-range forces were updated every step and the long-range forces were updated every three steps. The trajectories were saved at each 50.0 ps interval for analysis. The MD simulation results were analysed using the Simulation Interaction Diagram tool. The stability of MD simulations was monitored by Simulation Quality Analysis tool. The stability of the MD simulation results was confirmed by additional MD simulation runs.

### Microscale thermophoresis

METTL3/14 complex (26342, Cayman Chemical) was labelled via His-tag with Monolith His-Tag Labeling Kit RED-tris-NTA 2^nd^ Generation (NanoTemper Technologies GmbH). The labelled complex was diluted in MST buffer (10 mM Na-phosphate buffer, pH 7.4, 1 mM MgCl_2_, 3 mM KCl, 150 mM NaCl, 0.05% Tween-20) and centrifuged at 17 000×g for 10 min to avoid protein aggregates in MST. 20 nM labelled METTL3/14 complex was incubated with decreasing concentrations of SMT2457 for 30 min at room temperature. Samples were then loaded into Monolith Premium Capillaries (NanoTemper Technologies GmbH). Measurements were done with Monolith NT.115 instrument (NanoTemper Technologies GmbH) using red LED source, with power set at 100% and medium MST power at 25 °C. K_d_ values were calculated using the MO.Affinity Analysis software v2.3 (NanoTemper Technologies GmbH). Data are presented as ΔF_norm_ values ± SD from n = 4 independent experiments.

### Midbrain dopamine neurons and m6A regulator treatment

The midbrain floors were dissected from the ventral mesencephalic of 13 days old NMRI strain mouse embryos following the published procedure (L. Y. Yu et al., 2008). The tissues were incubated with 0.5% trypsin (103139, MP Biomedicals) in HBSS (Ca^2+^/Mg^2+^-free) (14170112, Invitrogen) for 20 min at 37 °C, then mechanically dissociated. Equal volumes of cell suspension were plated onto the centre of the 96-well plates coated with poly-L-ornithine (P- 8638, Sigma-Aldrich). The cells were grown in DMEM/F12 medium (21331–020, Invitrogen/Gibco) containing N2 supplement (17502–048, Invitrogen/Gibco), 33 mM D-glucose (G-8769, Sigma;), 0.5 mM L-glutamine (25030–032, Invitrogen/Gibco), and 100 μg/ml Primocin (ant-pm-05, Invivo Gen).

For survival assay, the cells were grown with different concentrations of METTL3/14 activators (10nM-1µM). Human recombinant GDNF (100 ng/mL) (P-103-100, Icosagen) or a condition without any trophic factor were used as positive and negative controls, respectively. After growing five days, the neuronal cultures were fixed and stained with anti-tyrosine hydroxylase (TH) antibody (MAB318, Millipore Bioscience Research Reagents). Images were acquired by ImageXpress Nano automated imaging system (Molecular Devices, LLC). Immunopositive neurons were counted by CellProfiler software, and the data was analysed by CellProfiler analyst software (McQuin et al., 2018). The results are expressed as % of cell survival compared to GDNF-maintained neurons (Mahato et al., 2020).

For rescue from neurotoxin treatment, the cells were grown for 5 days without any trophic factors in 96-well plates. Then, the cells were treated with 6-hydroxydopamine hydrochloride (6-OHDA, 15 µM, H4381, Sigma Aldrich) and METTL3/14 activators (100 nM or 1 µM). After three days the neuronal cultures were fixed and stained with anti-TH antibodies. Images were acquired by ImageXpress Nano high-content imaging equipment. Immunopositive neurons were counted by CellProfiler software, and the data was analysed by CellProfiler analyst software. The results are expressed as % of cell survival compared non-treated neurons.

All animal experiments were carried out following European Community guidelines for the use of experimental animals and approved by the Finnish National Experiment Board (License number: ESAVI/12830/2020) and by the Laboratory Animal Centre of the University of Helsinki (license no. KEK20-015; 2.7.2020).

### Human iPS-cell derived dopamine neurons and m6A regulator treatment

iCell® DopaNeurons (01279, FUJIFILM Cellular Dynamics) were seeded according to FUJIFILM Cellular Dynamics user protocol onto the 96-well plates coated with poly-L-ornithine (P3655, Sigma-Aldrich) and laminin (L2020, Sigma-Aldrich). Equal volumes of cell suspension were plated onto the centre of the dish. The cells were grown for 5 days in iCell Neural Base Medium 1+ iCell Neural Supplement B (M1010, M1029, FUJIFILM Cellular Dynamics). Then the cells were treated with 6-OHDA (50 µM) and different concentration of METTL3/14 activator (1 nM-1 µM), without any trophic factor or with GDNF (100 ng/ml) as negative and positive control, respectively. After three days, the cells were fixed and stained with anti-TH antibody. Images were acquired by ImageXpress Nano automated imaging system. Immunopositive neurons were counted by CellProfiler software, and the data was analysed by CellProfiler analyst software. The results are expressed as % of cell survival compared non-treated neurons.

### Measurement of m6A activator effects on human dopamine neuron neurite outgrowth

iCell® DopaNeurons were grown in 96-well plates with METTL3/14 activator (100 nM), or GDNF (100 ng/ml) or without any tropic factor as positive and negative control. After five days, the cells were fixed and stained with anti-Map2 antibody (AB5622, Sigma-Aldrich), anti-TH antibody and DAPI (D9542, Sigma-Aldrich). Images were acquired by ImageXpress Nano automated imaging system. Neurite outgrowth features of the neurons including total outgrowth, processes, and branches were assessed by CellProfiler software based on the immunocytochemistry of the neuronal marker Map2 (Pool et al. 2008). The total neurite length (µm) was measured based on Map2 positive area whereas neuronal cell numbers were counted based on the double-positive signal of Map2 and DAPI, and total neurite length per cell was calculated as the average of the total neurite length divided by the average of the total number of neuronal cells (Chang et al., 2019; Pansri et al., 2021).

### METTL3/14 activator C4 treatment in rat 6-OHDA model of Parkinson’s disease Experimental animals

Male Wistar rats (250–280 g) were housed in groups of 3 to 4 under a 12-h light-dark cycle at an ambient temperature of 20–23 °C. Food pellets (Harlan Teklad Global diet, Holland) and tap water were available ad libitum. During surgery days, the animals were housed separately for 24 h. All stereotaxic operations were performed under isoflurane anaesthesia as described in detail earlier (Voutilainen et al., 2009). Animals received 5 mg/ml of Carprofen (Rimadyl®, Zoetis Animal Health ApS, Denmark) and Buprenorphine 1 mg/ml (Temgesic, Schering-Plough) for post-operative pain relief. The experimental design was approved by the Committee for Animal Experiments of the University of Helsinki and the Chief veterinarian of the County Administrative Board. Behaviour of the animals was observed on daily basis and their weight was measured every week.

### Administration of 6-OHDA, GDNF and METTL3/14 activator C4

The design of the experiments is shown in Fig. 3a. Injections of 6-OHDA were done essentially as described earlier (Voutilainen et al., 2009), except the dosing into three injection sites. The animals received unilateral injections totalling 3×2 μg of 6-OHDA (Sigma Chemical CO, St. Louis, MO, USA; calculated as free base and dissolved in ice-cold saline with 0.02% ascorbic acid) in three deposits (2 μg/ 4 μl each) in the left striatum using coordinates relative to the bregma (A/P + 1.6, L/M + 2.8, D/V−6; A/P 0.0, L/M +4.1, D/V −5.5 and A/P −1.2, L/M +4.5, D/V −5.5) (Paxinos & Watson, 1997; Penttinen et al., 2016). Thus, the lesion used in the present study was different than in our earlier studies, where 20 μg of 6-OHDA was injected in one location in the neurorestoration paradigm (Lindholm et al., 2007; Voutilainen et al., 2009). The rats were divided into treatment groups according to their amphetamine-induced rotations at two weeks post-lesion. A brain infusion cannula was implanted to coordinates A/P + 1, L/M + 2.7, D/V −5 and secured to the skull using three stainless steel screws and polycarboxylate cement (Aqualox; VOCO, Germany). The tip of the infusion cannula was placed in the left striatum between the 6-OHDA injection sites. Cannula was connected via 2.5-cm-long catheter tubing to an osmotic pump (Alzet model 2002, Durect Corp., CA, USA), which was placed into a subcutaneous pocket between the shoulder blades.

The efficacy of METTL3/14 activator compound C4 was tested at two different doses by intracranial delivery and compared its effects to GDNF (Voutilainen et al., 2009). Two weeks after 6-OHDA treatment rats received the METTL3/14 activator C4 at two doses, PBS and GDNF intrastriatally using Alzet minipumps for four weeks. Four groups of rats were used: in one group PBS (11 rats) was used as vehicle control, the control group received GDNF (7 rats) at 3 μg/24 h and the two other groups received C4 at 0.3 μg/24 h or 1.5 μg/24 h (both 10 rats). After four weeks, Alzet minipumps were removed. Amphetamine-induced rotometry was carried out at weeks 2, 6, 9 and 12 (Voutilainen et al., 2011).

After behavioural tests, rats were perfused, and brains analysed for TH immunoreactivity in IHC (Voutilainen et al., 2009). TH-immunoreactive (TH-positive) cell bodies in the SNpc were counted using an automated deep convolutional neural networks (CNN) algorithm and cloud-embedded AiforiaTM platform. This computer-assisted cell counting method based on supervised machine learning and automated image recognition is described in detail and validated in (Penttinen et al., 2018). Optical density of the TH-positive fibres in the dorsal striatum was measured bilaterally from 3 different rostro-caudal levels through the striatum (approximately A/P +1.2 mm, +0.48 mm, and –0.26 mm relative to the bregma). The total magnitude of striatal denervation was analysed as an average reduction in the optical density at the three levels measured. Digital images of the TH-immunostained sections were acquired with Pannoramic P250 Flash II whole slide scanner (3DHistech) (Albert et al., 2021; Renko et al., 2021).

### Behavioural analysis

D-amphetamine-induced rotational behaviour was measured at weeks 2, 6, 9, and 12 post lesion in automatic rotometer bowls (Med Associates, Inc., Georgia, USA) as previously described (Lindholm et al., 2007; Ungerstedt and Arbuthnott, 1970). Following a habituation period of 30 min, a single dose of d-amphetamine (2.5 mg/kg, Division of Pharmaceutical Chemistry, University of Helsinki, Finland) was injected intraperitoneally (i.p.). The rotation sensor recorded complete (360^◦^) clockwise and counter clockwise uninterrupted turns for a period of two hours and ipsilateral rotations were assigned a positive value.

### Supplementary methods

### RNA extraction and RT-qPCR

Flow cytometry-sorted cells were lysed with TRI Reagent™ solution (AM9738, Invitrogen) and RNA isolated following manufacturer’s protocol. Synthesis of cDNA was carried out using oligo(dT) primers and Maxima H Minus Reverse Transcriptase (EP0753, Thermo Fisher Scientific). Relative gene expression was determined by using LightCycler 480 SYBR Green I Master (04887352001, Roche) and specific primers on a LightCycler® 480 Real-Time PCR System. Gene expression was measured in triplicates and normalized to the housekeeping gene actin.

### Neuronal culture and microinjections

Culture of mouse superior cervical ganglion (SCG) sympathetic neurons, dorsal root ganglion (DRG) sensory neurons, and microinjection of these neurons was performed as described earlier (Mätlik et al., 2015; L. Y. Yu et al., 2003). Briefly, the SCG neurons of postnatal day 1–2 NMRI strain mice or DRG neurons from E16 embryos were grown 6 DIV on poly-L-ornithine-laminin (P3655 and CC095, Sigma-Aldrich) coated dishes with 30 ng/ml of mouse NGF (G5141, Promega). Then the neurons were microinjected with METTL3/14 activators (200 nl) together with Dextran Texas Red (MW 70000 Da) (D1864, Invitrogen, Molecular Probes), that facilitates identification of successfully injected neurons, in PBS or PBS only (negative control). Recombinant MANF protein (P-101-100, Icosagen) in PBS at 200 ng/ul was used as positive control. Next day, tunicamycin (2 µM) (ab120296, Abcam) was added and living fluorescent (Dextran Texas Red-containing) neurons were counted three days later and expressed as percentage of initial living fluorescent neurons counted 2–3 hours after microinjection.

### Immunocytochemistry

SCG or DRG neurons were cultured on 96-well plates for 6 days. Then the cells were fixed with 4% PFA and stained with the following antibodies (1: 500 dilutions used for all): rabbit anti-m6A (202 003, Synaptic Systems), rabbit anti-METTL3 (15073-1-AP, Proteintech), mouse anti-Tau1 (MAB3420, Sigma-Aldrich). The nuclei were stained with DAPI (D9542, Sigma-Aldrich). Images were acquired by ImageXpress Nano automated imaging system.

