## Supplementary File S1 for "Small molecule activation of m6A mRNA methylation as a novel approach for neuroprotection"

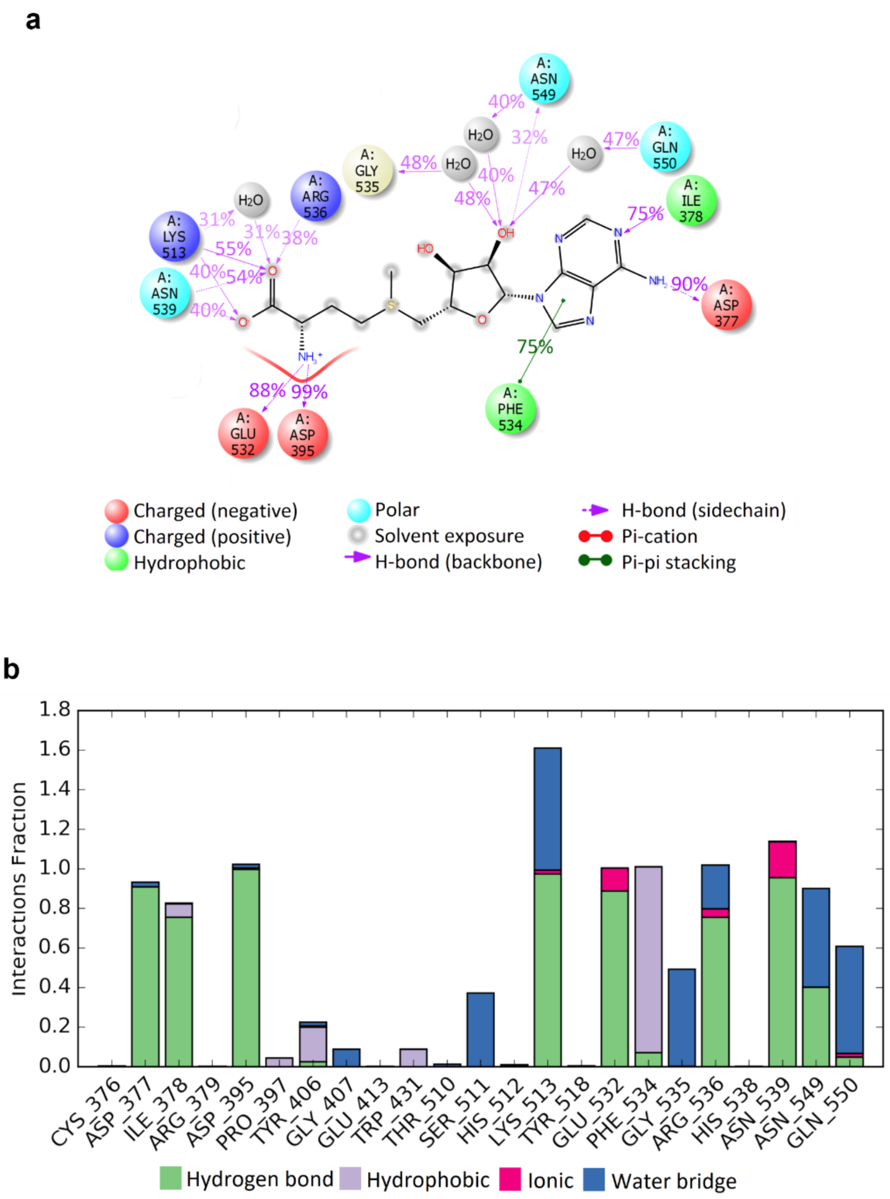


**Suppl. Fig. S1: a,** 2D summary diagram of the 50 ns molecular dynamics calculated contacts between METTL3/METTL14 and SAM. Interactions that occur more than 30% of the simulation time are shown. **b,** Molecular dynamics calculated contacts for the METTL3/METTL14/SAM complex (PDB ID: 5K7W). The stacked bar charts are normalized over the course of the trajectory. Values over 1.0 are possible as some protein residue may make multiple contacts of same subtype with the ligand.

**a**

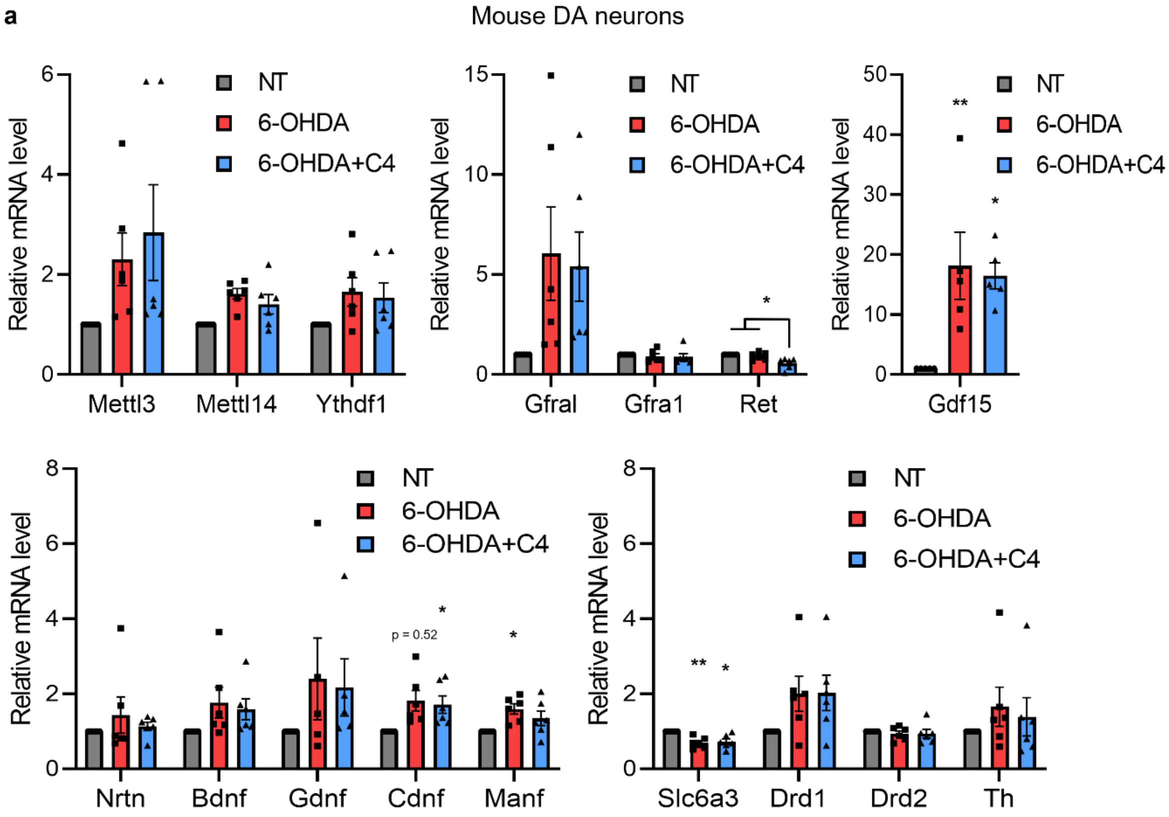

Mouse DA neurons
Relative mRNA level
NT
6-OHDA
6-OHDA+C4
Mettl3
Mettl14
Ythdf1
Gfral
Gfra1
Ret
Gdf15
Nrtn
Bdnf
Gdnf
Cdnf
Manf
p = 0.52
Slc6a3
Drd1
Drd2
Th

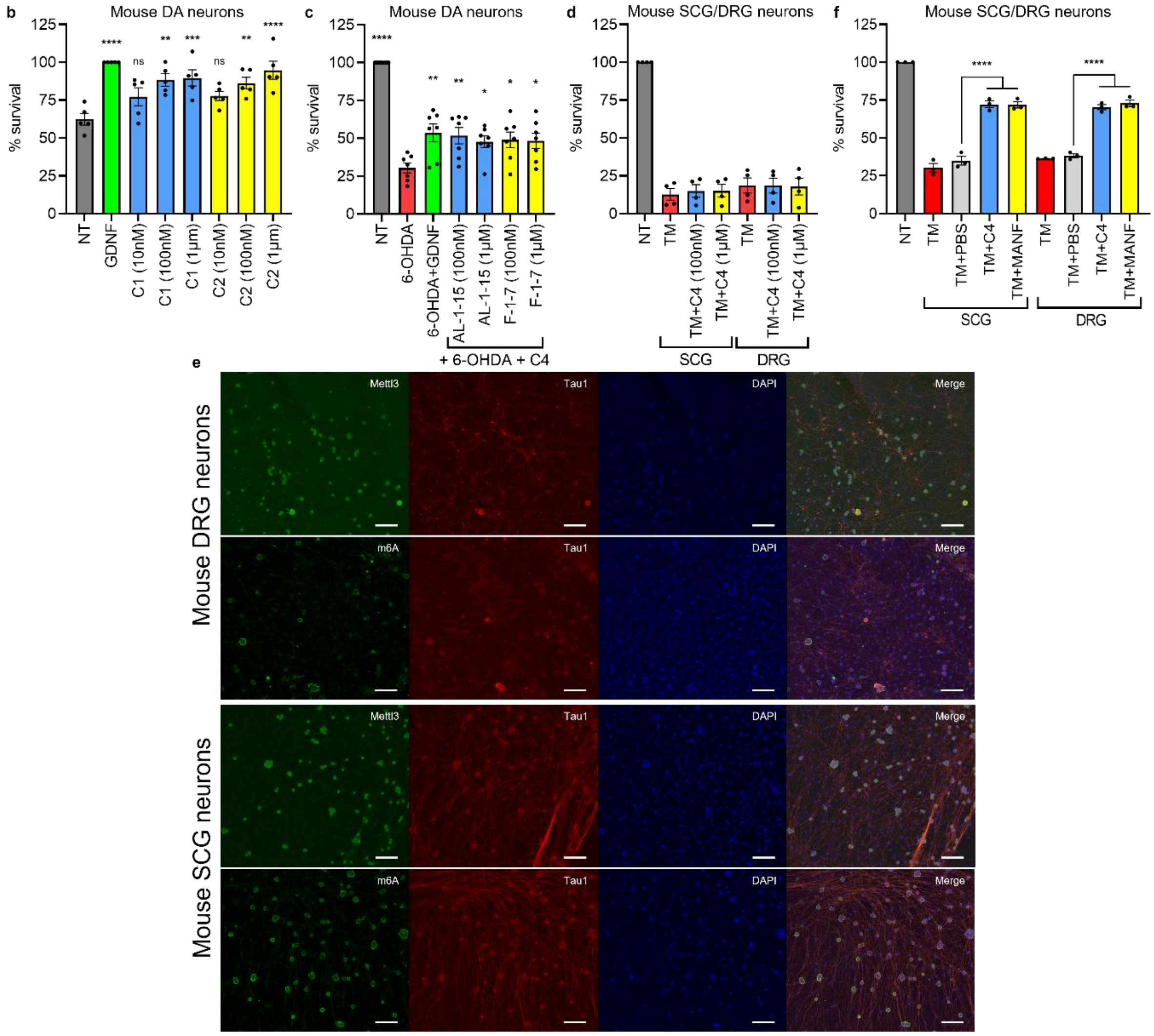


**Suppl. Fig. S2: a,** Gene expression profile of flow cytometry-sorted TH+ mouse dopamine neurons. Ythdf1 – YTH N6-methyladenosine RNA binding protein F1, Gfral – Gdnf family receptor alpha like, Gfra1 – Gdnf family receptor alpha 1, Ret – Ret proto-oncogene, GDF15 – Growth differentiation factor 15, Nrtn – Neurturin, Bdnf – Brain derived neurotrophic factor, Gdnf – Glial cell line-derived neurotrophic factor, Cdnf – Cerebral dopamine neurotrophic factor, Manf – Mesencephalic astrocyte-derived neurotrophic factor, Slc6a3 – Solute Carrier Family 6 Member 3 (dopamine transporter), Drd1/2 – Dopamine receptor D1/2, Th – tyrosine hydroxylase. **b,** Survival of mouse DA neurons in growth factor deprivation assay with C1 or C2 compared to GDNF (100ng/ml). **c,** Survival of mouse DA neurons with GDNF (100ng/ml) or combination of C4 (100nM) and AL-1-15 or F-1-7 in the presence of 6-OHDA (15μM). **d,** Survival of SCG and DRG neurons in the presence of TM (2μM) and C4 in media. **e,** Representative images of mouse DRG and SCG neurons stained with Mettl3 (green), m6A (green), Tau1 (red) antibodies and counterstained with DAPI (blue), scale bar 200μm. **f,** Survival of SCG and DRG neurons in the presence of TM (2μM) and microinjected with C4 or MANF. *p<0.05, **p<0.01, ***p<0,001, ****p<0.0001
